# Stearoyl coenzyme A desaturase 1 (SCD1) regulates foot-and-mouth disease virus replication by modulating host cell lipid metabolism and 2C-mediated replication complex formation

**DOI:** 10.1101/2024.05.23.595652

**Authors:** Bonan Lv, Yuncong Yuan, Zhuang Yang, Xingran Wang, Jianjun Hu, Yidan Sun, Hang Du, Xuemei Liu, Huimin Duan, Ruyi Ding, Zishu Pan, Xiao-Feng Tang, Chao Shen

**Affiliations:** Hubei Key Laboratory of Cell Homeostasis, College of Life Sciences, Wuhan University, Wuhan 430072, China; State Key Laboratory of Virology, College of Life Sciences, Wuhan University, Wuhan 430072, China

**Keywords:** foot-and-mouth disease virus, SCD1, host lipid metabolism, oleic acid, replication complex

## Abstract

The life cycle of foot-and-mouth disease virus (FMDV) is tightly regulated by host cell lipid metabolism. In a previous study, we successfully established a BHK-21 cell model (BHK-Op) for persistent FMDV infection by single-cell clone selection and isolated a virus-negative cell line (BHK-VEC cells) from high-passage BHK-Op cells. BHK-VEC cells could be not infected with FMDV. By conforming the transcriptome data of BHK-VEC cells and BHK-21 cells, we identified that the stearoyl coenzyme A desaturase 1 (SCD1), a key enzyme for the fatty acid metabolism, regulates FMDV replication. SCD1 overexpression or exogenous addition of oleic acid (OA), a product of SCD1 enzyme activity, promoted or upregulated FMDV replication in BHK-21 cells or SCD1 knockdown cells, respectively. Interestingly, overexpression of SCD1 or exogenous addition of OA restored the FMDV infection and replication in BHK-VEC cells. Exogenous addition of OA also promoted FMDV replication in BHK-Op. SCD1 recruited the nonstructural FMDV protein 2C to the detergent-resistant membrane located in the perinuclear nucleus to form replication complex. Inhibition of SCD1 enzyme activity resulted in significantly decreased number of FMDV replication complexes with also abnormal morphology. Importantly, inhibition of SCD1 enzyme activity effectively suppressed replication of other positive-sense RNA viruses, such as REO176, PV1 and EV71. Our results demonstrated that, as a key host regulator of RNA virus replication, SCD1 is a potential target for developing novel drugs against positive-sense RNA virus infection.

## Introduction

Foot-and-mouth disease (FMD) is an acute, highly contagious viral disease caused by foot-and-mouth disease virus (FMDV), which infects even-hoofed animals (such as cattle, sheep, and goats) causing rapid disease onset and high mortality. FMDV is a positive-strand RNA virus of the family *Picornaviridae* and genus *Aphthovirus*. Recent studies have demonstrated that many positive-strand RNA virus infections are accompanied by changes in host intracellular lipid metabolism, including remodeling of the host cell endosomal system, decreased fatty acid oxidation, increased *ab initio* lipid synthesis and fatty acid transport rate^[1]^. Positive-strand RNA viruses display increased survival and replication efficacies in host cells by coordinating virus– host interactions to remodel host cell membranes and lipid metabolism and altering the microenvironment^[2]^. Dengue virus (DENV), a *Flavivirus* member, inhibits the *ab initio* synthesis of lipids and disrupts the formation of transporter vesicles, which enhances viral infection^[3]^. In DENV-infected cells, fatty acid synthase (FASN) is relocated to the DENV replication site where viral nonstructural protein 3 (NS3) colocalizes and interacts with FASN. Purified recombinant NS3 stimulates the enzymatic activity of FASN *in vitro*, and inhibition of FASN enzymatic activity inhibits DENV infection, indicating that DENV relies on the host cell fatty acid biosynthesis pathway to establish its replication complex^[4–6]^. In cells infected with DENV, the rate of fatty acid synthesis was significantly increased compared to that in uninfected cells, and inhibition of lipid synthesis resulted in significantly decreased DENV replication^[4–6]^. Stearoyl coenzyme A desaturase 1 (SCD1) catalyzes the biosynthesis of monounsaturated fatty acids from saturated fatty acid precursors, and both the gene expression and enzyme activity of SCD1 were significantly upregulated early in cells infected with DENV2. This finding suggested that the catalytic products of SCD1 play important roles in the life cycle of this virus.

In FMDV-infected cells, the endoplasmic reticulum membrane is remodeled, and many new vesicular structures appear in the cytoplasm. The formation of these vesicular structures requires the participation of a large number of free fatty acids. The vesicular structures also contain nonstructural proteins of FMDV origin and cytokines of host cell origin^[7, 8]^. This suggests that host cell lipid metabolism is closely related to replication complex formation. In cells infected with several small RNA viruses, type III phosphatidylinositol 4 kinase catalyzes the substrate to produce high levels of PI4P, which is subsequently converted to cholesterol in the replication complex^[9]^. However, the role of host factors involved in lipid metabolism-mediated regulation of FMDV replication or replication complex formation has not been reported. Studies on the functions of *SCD1*–*4*, members of the SCD family have focused mainly on SCD1. *SCD1*–*3* are present in the Syrian hamster.^[10]^ SCD1 is a delta-9 desaturase and a rate-limiting enzyme in the biosynthesis of monounsaturated fatty acids from saturated fatty acid precursors in organisms. The enzymatic products of SCD1 (e.g., oleic acid (OA), diacylglycerol, phospholipids, triglycerides, palmitoleic acid, and cholesterol esters) are involved in the formation of the hepatitis C virus (HCV) replication complex^[11]^. Based on these observations, we speculated that SCD1 enzymatic activity may be crucial for FMDV replication. SCD1 contains four transmembrane helices (TM1– 4) that form the stem of the mushroom-shaped protein and a cytosolic structural domain that forms the top of the mushroom. TM2 and TM4, which are longer than TM1 and TM3, protrude into the cytosolic region, providing three of the nine histidine residues that coordinate two metal ions^[12, 13]^. OA increases fluidity and structural changes in the phospholipid membrane bilayer, including negative curvature of biological membranes, which is essential for the formation of closed vesicle-like structures in cells. SCD1 is the key enzyme in intracellular OA synthesis, and a significant decrease in the number of intracellular lipid droplets and specialized double-membrane vesicles was observed via electron microscopy after treating Huh7 cells with an SCD1 inhibitor^[14]^. However, the effect of OA metabolism on viral replication in host cells has rarely been investigated.

Here, we described an in depth investigation of the role of SCD1-regulated lipid metabolism in FMDV replication and the underlying molecular mechanisms. We found that SCD1 regulates FMDV replication by modulating host cell lipid metabolism and promoting 2C-mediated viral replication complex formation. In addition our study found that in SCD1 also regulates replication of justice RAN viruses such as REO176, PV1, EV71. Our findings provide new ideas for innovative anti-justice RNA virus strategies.

## Significance

Many positive-stranded RNA virus infections are accompanied by alterations in host cell intracellular lipid metabolism. Positive-stranded RNA viruses reshape host membranes and lipid metabolism by coordinating virus-host interactions to create a suitable microenvironment for survival and replication in host cells. In FMDV-infected host cells, the endoplasmic reticulum membrane is remodeled and many nascent vesicular structures appear in the cytoplasm. The formation of these vesicular structures requires the involvement of large amounts of free fatty acids, and the vesicular structures also contain a number of FMDV-derived nonstructural proteins and host cell-derived cytokines, suggesting that host cell lipid metabolism and the formation of the FMDV replication complex are closely related. 2C is a common viral protein in the replication complex of viruses of the family Picornaviridae and is thought to be an important component of cell membrane rearrangement and viral replication complex formation. SCD1 is an important protein involved in host cell lipid metabolism. We found that FMDV infection of host cells can lead to differential expression of SCD1, and inhibition of SCD1 expression or enzymatic activity can significantly inhibit FMDV replication, while FMDV replication was regulated when exogenous addition of SCD1 active product OA. Furthermore, we found that SCD1 is present in the FMDV replication complex and is involved in recruiting 2C to the detergent-resistant membrane. This suggested that lipid metabolites involved in FMDV replication complex formation play a key role in the regulation of viral replication and persistent infection. Our study will reveal the role of SCD1 in the lipid metabolism pathway and the molecular mechanism mediating the formation of the FMDV replication complex and thus regulating viral replication, explore the regulatory relationship between cellular lipid metabolism and FMDV replication and persistent infection, and provide a theoretical basis for an in-depth understanding of the pathogenesis of FMD and innovative antiviral strategies, providing a potential drug target against FMD virus.

## Methods

### Cell lines and virus

FMDV type O (Akesu/58/2002) was kindly provided by the Lanzhou Veterinary Research Institute, Chinese Academy of Agricultural Sciences. The FMDV titer was calculated using a plaque assay. The golden hamster kidney fibroblast line BHK-21 was obtained from the China Center for Type Culture Collection (CCTCC). PK-15 and Vero cells were donated to CCTCC. REO176, EV71 and PV1 were donated to the CCTCC.

### Cell culture

BHK-21 cells, BHK-Op cells (BHK-21 cells with persistent FMDV infection), and BHK-VEC cells (virus-negative cells isolated from among BHK-Op cells) were cultured in T25 culture flasks containing 4–6 mL of minimum essential medium (MEM) containing 10% fetal bovine serum (FBS; Every green) and 1% streptomycin and penicillin. All cells were cultured at 37°C with 5% CO_2_. When the cells reached 80–90% confluence, they were passaged, the old medium was aspirated and discarded, the cells were washed 2–3 times with phosphate-buffered saline (PBS), and the cells were digested using 1 mL of 0.5% trypsin solution. The trypsin digestion was stopped by adding two volumes of serum-containing medium to the trypsin-containing solution, after which the mixture was centrifuged at 1000 rpm for 5 min. The cells were resuspended in medium at a ratio of 1:4, after which the cells were supplemented with MEM and incubated at 37°C under 5% CO_2_.

The SCD1-specific inhibitor was dissolved in DMSO or and stored at −20°C. The fatty acids used included oleic acid (OA; 18:1), stearic acid (SA; 18:0), and palmitoleic acid (POA; 16:0). For example, cells were cultured in 12-well plates, and once they reached 60%–70% confluence, they were infected with FMDV (50% tissue culture infectious dose (TCID_50_) = 1×10^−4^/mL); 1 h later, the medium was replaced with MEM containing 2% FBS, followed by the addition of fatty acids or the indicated concentration of SCD1 inhibitor. The cells were collected at 16 or 24 h postinfection (hpi), and the RNA and protein levels were analyzed as previously described.

### Plasmids and DNA transfection

To construct the overexpression vector we first isolated total RNA from BHK-21 cells and then reverse transcribed it to create cDNA. The cDNA was then used as a template for amplification of the desired gene fragment using Phanta Max Super-Fidelity DNA Polymerase (Vazyme). The amplified target fragment was subcloned and inserted into the pHAGE-CMV-MCS-IZsGreen vector using the ClonExpress II One Step Cloning Kit (Vazyme). The successful construction of the plasmid was verified by enzymatic digestion and DNA sequencing.

The knockdown plasmid was constructed by first designing and synthesizing a 21-polymer oligonucleotide and annealing it to form a double stranded short interfering RNA (siRNA). The annealed double-stranded oligonucleotides were cloned between the *Hin*dIII and *Bgl*II restriction sites of the pSUPER.retro.puro plasmid (Oligoengine). The successful construction of the knockdown plasmid was verified by digestion using the restriction endonucleases *Eco*RI and *Hin*dIII.

Cells were cultured in a 12-well plate at 37°C, and transfection reagents A (1.6 μg of plasmid + 100 μL of opti-MEM per well) and B (1.6 μL of Lipofectamine 2000 + 100 μL of opti-MEM per well) were prepared when the cell density reached approximately 60%. Liquids A and B were mixed and allowed to stand for 15 min. The medium in the 12-well plate was discarded, the cells were washed 3 times with PBS, and the mixture of transfection reagents A and B was added. Then, the cells were incubated at 37°C with 5% CO_2_ for 4–6 h. The transfection medium was aspirated from the 12-well plate and discarded and the cells were incubated for 24 h in MEM supplemented with 2% FBS.

### Cytotoxicity assays

The cells were cultured in 96-well plates, and virus were added when the cell confluence was approximately 60%. Eight concentrations of each drug to be tested (diluted with DMSO) were selected, and eight replicates were used for each concentration. The treated 96-well plates were incubated at 37°C with 5% CO_2_. After 24 h, CCK-8 (Cell Counting Kit-8) reagent (Yeasen Biotech) was added (10% vol:vol), and the plates were subsequently incubated at 37°C with 5% CO_2_ for 2 h. Absorbance values were measured at 450 nm using an enzyme marker (Thermo Fisher). The cell survival rate was calculated as follows: cell survival rate = (absorbance value of experimental group − absorbance value of blank control group)/(absorbance value of solvent group − absorbance value of blank control group).

### RNA extraction and reverse transcription quantitative PCR (RT–qPCR)

The medium was removed from a 12-well plate, and 500 μL of RNAiso Plus was added to each well. Then, the mixture was incubated at 4°C for 30 min. Cells from each sample were extracted with 100 μL of chloroform and collected into RNase-free Eppendorf (EP) tubes, mixed thoroughly, incubated for 5 min, and centrifuged at 12000 × *g* at 4°C for 15 min. The supernatant was aspirated and transferred to a fresh RNase-free EP tube, mixed with an equal volume of isopropanol, incubated at 4°C for 20 min, and centrifuged at 12000 × *g* at 4°C for 15 min. The supernatant was discarded, and the precipitate was washed with 75% ethanol and centrifuged at 12000 × *g* at 4°C for 5 min. The supernatant was discarded, the precipitate was thoroughly dried, and the RNA was dissolved in prewarmed RNase-free water.

cDNA was synthesized by reverse transcription of the RNA using a Hifair II 1^st^ Strand cDNA Synthesis Kit (Yeasen), followed by chain-specific RT‒qPCR using SYBR Green dye and a CFX96 real-time PCR detection system. The expression level of *GAPDH* was used as an internal reference, and relative changes in the expression of target genes were calculated via the 2^−△△CT^method.

### Western blotting

Sodium dodecyl sulfate (SDS) loading buffer (containing RIPA strong lysis buffer) was added to a 12-well plate to lyse the cells. The protein lysate was collected in an EP tube, heated at 100°C for 10 min in a dry heat block, and subsequently centrifuged at 12000 rpm for 5 min. Samples were separated via 10% or 15% SDS‒polyacrylamide gel electrophoresis and subsequently transferred to polyvinylidene fluoride membranes. The membranes were blocked with 5% skim milk for 1–2 h and then incubated with a specific primary antibody (diluted with antibody diluent) at 4°C overnight, followed by washing three times with TBST and incubating with a secondary antibody (horseradish peroxidase-conjugated goat anti-rabbit or goat anti-mouse antibody) for 2 h at room temperature; the proteins were subsequently visualized with an enhanced chemiluminescence reagent. Antibodies used in this experiment will be listed in Table 1.

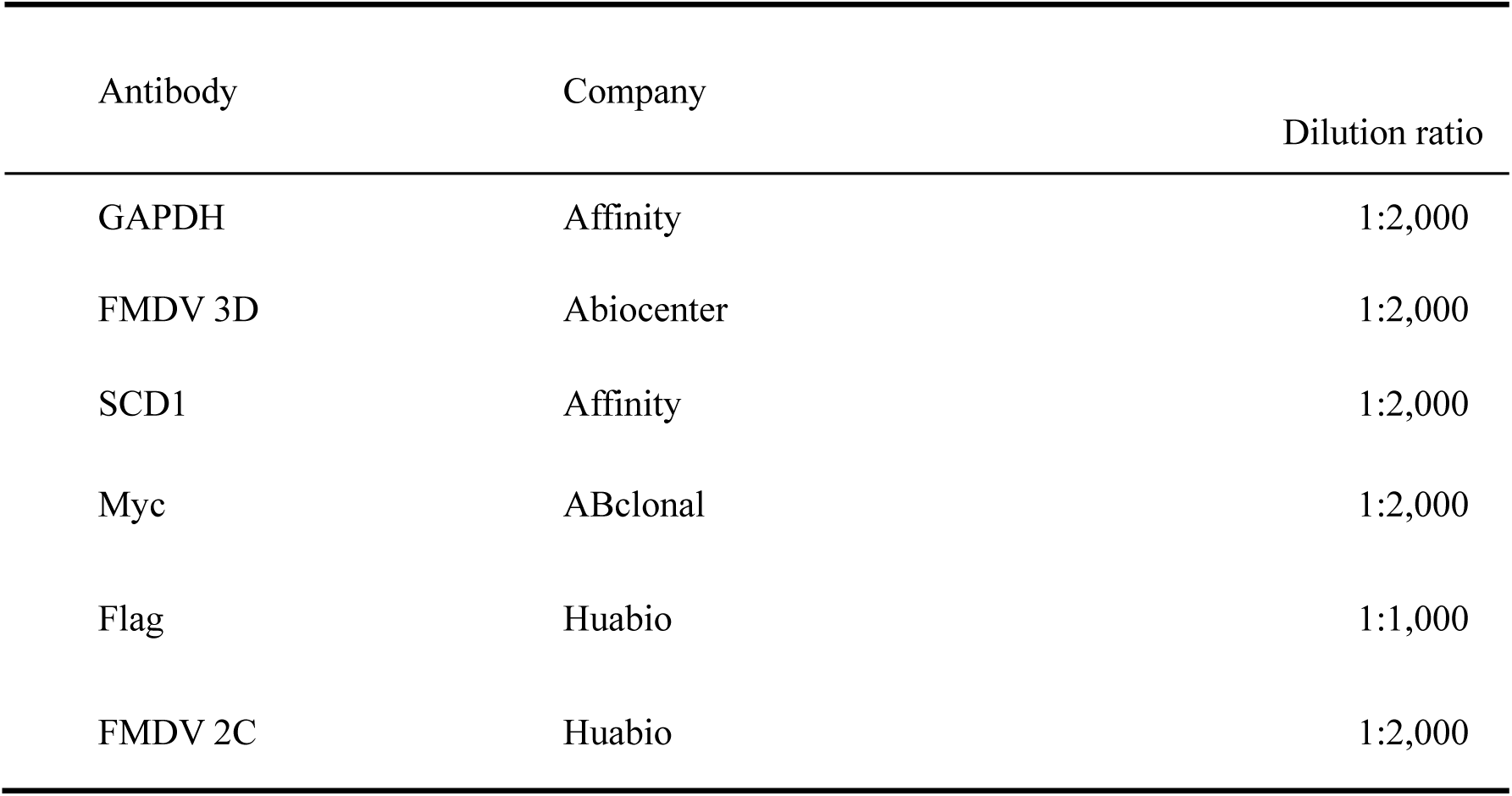
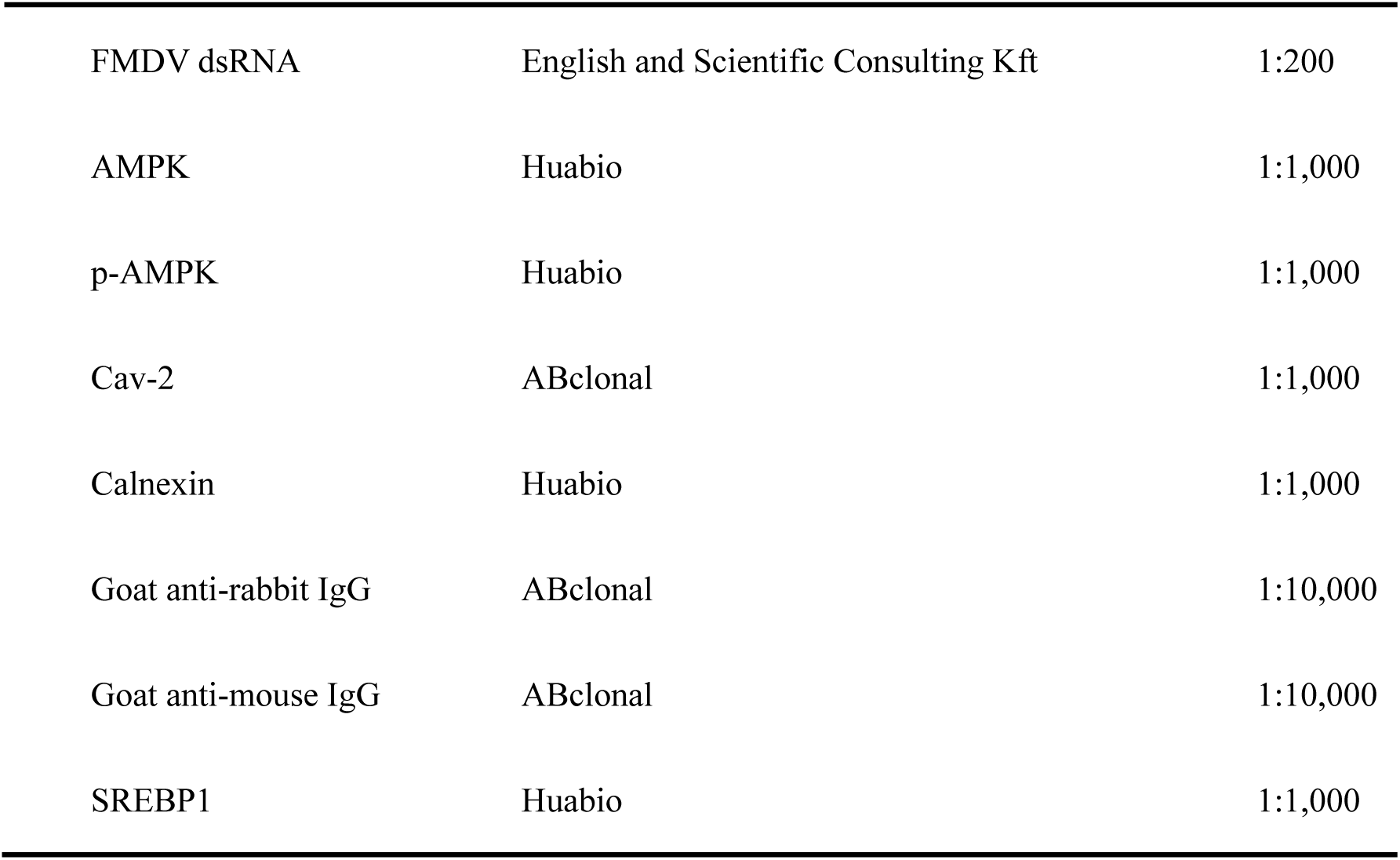
Antibodies used in this study.

### Viral titer assay

When the confluence of the BHK-21 cells in the T25 culture flasks reached 80–90%, the cells were digested with trypsin for passaging in 96-well plates. The viral stock solution to be tested was diluted with a gradient series of serum-free MEM. Diluted viral solution (100 μL) was added to each well (eight replicate wells for each concentration), and in the control group, the same volume of virus-free medium was added. Maintenance medium (100 μL) containing 2% FBS was added to each well, and the 96-well plate was incubated at 37°C with 5% CO_2_. Every 24 h, the cells in each well was observed under a microscope for cytopathic effects (CPE), and the number of wells in which CPE was observed at each dilution was counted for 3 days.

### Coimmunoprecipitation (Co-IP)

Plasmids were transfected into cells cultured in T25 flasks, and the cells were infected with FMDV (TCD_50_ = 1×10^−4^/mL) after 24 h. After 1 h of treatment, the culture medium was changed to MEM containing 2% FBS, and the incubation was continued for 16 h. Each flask was aspirated and washed twice with PBS and 1 mL of IP lysis mixture (with protease inhibitor cocktail and phenylmethylsulfonyl fluoride preadded); then, the flasks were incubated for 10 min at 4°C in a shaker. The samples were collected into EP tubes using a cell scraper. The antibodies were added to magnetic beads treated with binding/washing buffer (diluted 1:2000 with binding/washing buffer) and incubated overnight on a rotating mixer at 4°C. The samples (400 μL) were then incubated for 2 h on an rotating mixer at 4°C. Magnetic separation was performed on a magnetic holder, and the samples were washed four times with binding/washing buffer. Finally, immunoprecipitated protein samples were prepared for subsequent immunoblotting using a denaturing elution method.

### Luciferase reporter assay

The promoter region of the *SCD1* target gene (1000 bp upstream and downstream of the start codon, containing the classical serum response element (SRE)) was subcloned and inserted into a reporter vector containing the luciferase gene (pGL3-basic^[15]^). The constructed pGL3-SRE reporter plasmid (100 ng) was cotransfected with phRL-TK (2 ng; internal reference, containing the sea kidney luciferase gene) into BHK-21 cells cultured in a 24-well plate for 24 h. Then, the 1× passive lysis buffer was added and the cells were completely lysed on a shaker for 20–30 min before collection. The intensity of the dual-luciferase fluorescence signal was detected using the Promega GLOMAX operating system^[16]^.

### Immunofluorescence (IF) assay

Cells were cultured in 24-well plates, and once the cells reached 60%–70% confluence, they were fixed with 4% paraformaldehyde for 30 min. After fixation, the cells were washed three times with sterile PBS, permeabilized with 0.5% Triton X-100 (diluted in PBS) for 12 min, washed three more times with sterile PBS, and blocked with 5% BSA (diluted in PBS) for 30 min after which the blocking solution was aspirated and discarded. Primary antibody (diluted with 1% BSA) was added to the cells, which were then incubated overnight at 4°C. The cells were washed three times with sterile PBST, the secondary antibody dilution was added, and the mixture was incubated at room temperature for 2 h. At the end of the incubation, the cells were washed three times with sterile PBST and observed under a fluorescence microscope^[17]^.

### Transmission electron microscopy (TEM)

Approximately one-third of the cell suspension was mixed with a 1:1 mixture of 0.1 M phosphate buffer and 2.5% glutaraldehyde. The mixture was allowed to stand for 10 min and then centrifuged at 1000 rpm for 10 min. The supernatant was aspirated and discarded. Glutaraldehyde (2.5%) was added for approximately 20 min to fix the cells, after which the samples were centrifuged at 2000 rpm for 10 min and stored in a refrigerator at 2–8 °C. Afterward, the samples were vacuumed for 2–3 h using a vacuum device, and the cells were fixed for a second time. After centrifugation to remove the supernatant, the degree of cell aggregation was observed. If the sample cells had not formed clumps or the observed clumps were small, 0.6%–1% agar was added, and aggregates were cut into suitably sized clumps; if the cells had formed larger clumps, they were cut into suitably sized clumps directly. The cut clumps were washed at least twice with 0.1 M phosphate buffer for approximately 10 min each time (or mixed with 0.1 M phosphate buffer, stored overnight in a refrigerator at 2–8 °C and washed at least once the next day). The clumps were then fixed with 1% osmium tetroxide for 1–2 h at room temperature in the dark, and then fixed for observation via TEM ^[18]^.

### Detergent-soluble and detergent-resistant membranes preparation and flotation assay

The cells were lysed in TNE buffer (150 mM NaCl, 1.0% Triton X-100, 3 mM EDTA, and 20 mM Tris-HCl), and a mixture of phosphatase and protease inhibitors was added. The cell lysate was then homogenized 10 times by passage through a 23-gauge needle and incubated on ice for 1h. The lysate was centrifuged at 14,000 × *g* at 4°C for 30 min, and the supernatant was collected as the detergent-soluble fraction. The remaining (precipitated) fraction was resuspended in the same lysis buffer supplemented with 0.5% SDS and 2 mM DTT and subjected to brief sonication, following which the supernatant was collected as the detergent-resistant fraction^[19]^.

### Statistical analysis

Student’s *t* test was used for statistical analysis with GraphPad Prism version 8.0, ImageJ, and Image Lab 5.2 software. A *P* value < 0.05 was considered to indicate statistical significance. The data are expressed as the mean ± standard deviation. *, *P* < 0.05; **, *P* < 0.01; ***, *P* < 0.001; ns, not significant.

## Results

### SCD1 is essential host factor for regulating FMDV replication

We previously established the persistently FMDV infected BHK cell line (BHK-Op) by single-cell selection^[20]^, and an FMDV-negative cell clone (BHK-VEC) was isolated by single-cell clonal screening of high-passage BHK-Op cells (detoxified cells)^[21]^. To understand the reason for the virus resistance of BHK-VEC cells, we analyzed the differential genes through transcriptome analysis between BHK-VEC cells and BHK-21 cells. We identified differentially expressed genes (DEGs) between BHK-21 and BHK-VEC cells using transcriptome sequencing. These DEGs were related to fatty acid metabolism pathways, and fifteen representative DEGs were selected for validation by RT‒qPCR (Figure S1A). *SCD1* is one of 15 DEGs. *SCD1* mRNA levels were significantly downregulated in BHK-VEC cells and significantly upregulated in BHK-21 cells infected with FMDV compared with uninfected BHK-21 cells. SCD1 expression in BHK-Op cells was significantly increased wih BHK-21 cells (Figure 1A). The decreased *SCD1* expression level in BHK-VEC cells may be related to the absence of FMDV in these cells.

**Figure 1.**
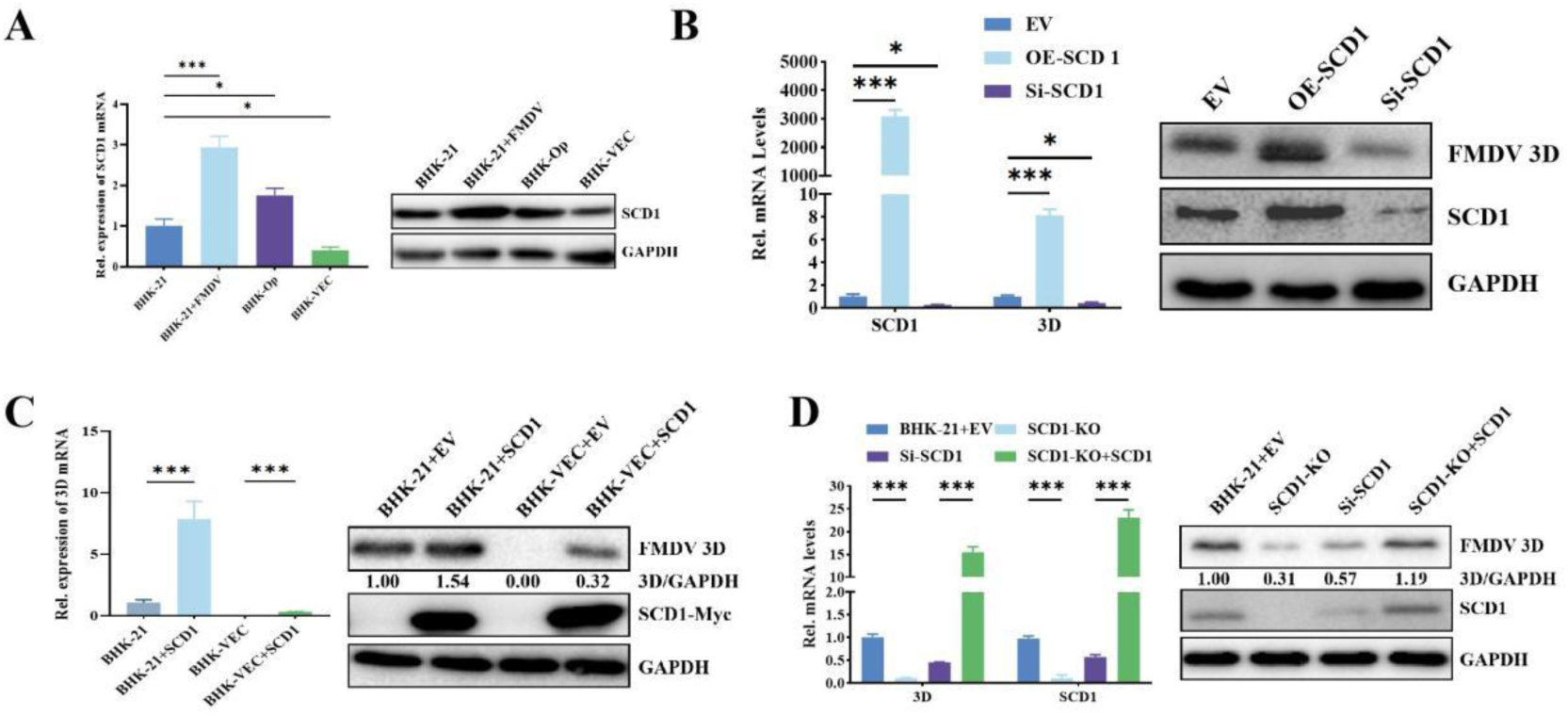
Identification of the function of SCD1 in regulating foot-and-mouth disease virus (FMDV) replication. (A) mRNA levels of SCD1 mRNA levels in BHK-21 cells, BHK-21 cells infected with FMDV, BHK-VEC cells and BHK-Op cells; (B) BHK-21 cells and BHK-VEC cells were transfected with a phage-CMV-IZsGreen (EV) SCD1-overexpression (OE-*SCD1*) plasmid or RNA interference plasmid (Si-*SCD1*) for 24 h and were then infected with FMDV for 16 h. Detection of 3D and SCD1 expression in FMDV by qPCR and western blotting; (C) BHK-VEC cells were transfected with the OE-*SCD1* plasmid for 24 h and then infected with FMDV for 16 h. Detection of 3D and SCD1 expression in FMDV by qPCR and western blotting; (D) mRNA and protein levels of SCD1 in the *SCD1*-KO cell line and the mRNA and protein concentrations of 3D were collected 16 h after FMDV infection. The protein bands were quantified in grayscale by using Image-Pro Plus 6.0 software for the western blot. *n* = 3 for each group of experiments, and three parallel samples were combined for western blotting. \**P* < 0.05, n.s., not significant.

To investigate the functions of the SCD family, we constructed RNAi plasmids targeting homologous regions in the *SCD1*, *SCD2*, and *SCD3* genes. These RNAis effectively suppressed *SCD1*, *SCD2*, and *SCD3* expression, with the most significant inhibition observed for *SCD1* expression (Figure S1B&C). Inhibition of *SCD1* expression in FMDV-infected BHK-21 cells resulted in a significant decrease in the mRNA levels of FMDV 3D, an RNA polymerase encoded by FMDV (Figure S1B & C). Consistent with the mRNA level, the FMDV 3D protein level was also significantly decreased in FMDV infected cells following *SCD1* inhibition (Figure S1B & C). These results suggested that *SCD1* modulates FMDV replication in infected BHK-21 cells.

To investigate the regulatory effect of *SCD1* gene expression on FMDV replication, we overexpressed and knocked down *SCD1* in BHK-21 cells. At 16 h posttransfection (hpt) with overexpression or knockdown constructs, the transfected cells were infected with FMDV. FMDV replication was significantly upregulated in *SCD1-*overexpressing cells and significantly downregulated in cells with *SCD1* knockdown (Figure 1B). Overexpression of SCD1 promoted FMDV replication (Figure 1B). Surprisingly, overexpression of SCD1 in BHK-VEC cells restored FMDV replication (Figure 1C). Our results demonstrated that SCD1 is an important host factor necessary for FMDV replication.

We applied CRISPR technology to construct an *SCD1*-knockout (KO) BHK-21 cell line. *SCD1* knockout significantly decreased the 3D protein level in *SCD1*-KO cells compared with BHK-21 cells (Figure 1D), suggesting that the replication of FMDV was dependent on SCD1 expression. Subsequently, we overexpressed *SCD1* in *SCD1*-KO cells (Figure 1D) and could restore 3D expression. These results demonstrated that SCD1 plays a critical role in regulating FMDV replication.

### SCD1 enzyme activity is essential for upregulating FMDV replication

To investigate the role of SCD1 enzyme activity in regulating FMDV infection, we treated BHK-21 cells with MK8245, an inhibitor of SCD1 enzyme activity. We first assayed the cytotoxicity of MK8245 toward BHK-21 cells using a CCK-8 assay; and the IC50 value of MK8245 was ≥50 μM. FMDV replication was significantly downregulated by MK8245 in a dose-dependent manner, and FMDV 3D protein expression was also significantly decreased (Figure 2A&B). The addition of MK8245 significantly decreased the titer of FMDV (Figure 2C). To elucidate the role of the SCD1 enzymatic product (unsaturated fatty acids) in FMDV replication, we evaluated the relationship between FMDV replication and unsaturated fatty acids by adding exogenous stearic acid (SA), a substrate of SCD1. The ability of SA to upregulate FMDV 3D protein and RNA levels was further enhanced upon *SCD1* overexpression (Figure 2D). SA strongly promotes FMDV replication, suggesting that not only the SCD1-mediated conversion of SA to OA but also other unidentified host cell lipid metabolism pathways are important for FMDV replication. In BHK-21 cells, exogenous SA significantly upregulated FMDV replication (Figure 2E), suggesting that SCD1 can still catalyze the conversion of SA into unsaturated fatty acids. In *SCD1*-KO cells, exogenous SA did not affect FMDV replication (Figure 2F), suggesting that the SCD1-meidated enzymatic conversion of SA into OA, not SA itself, promoted FMDV replication. These data demonstrated that monounsaturated fatty acids, especially OA, the product of SCD1 enzymatic activity, are specifically required for FMDV replication.

**Figure 2.**
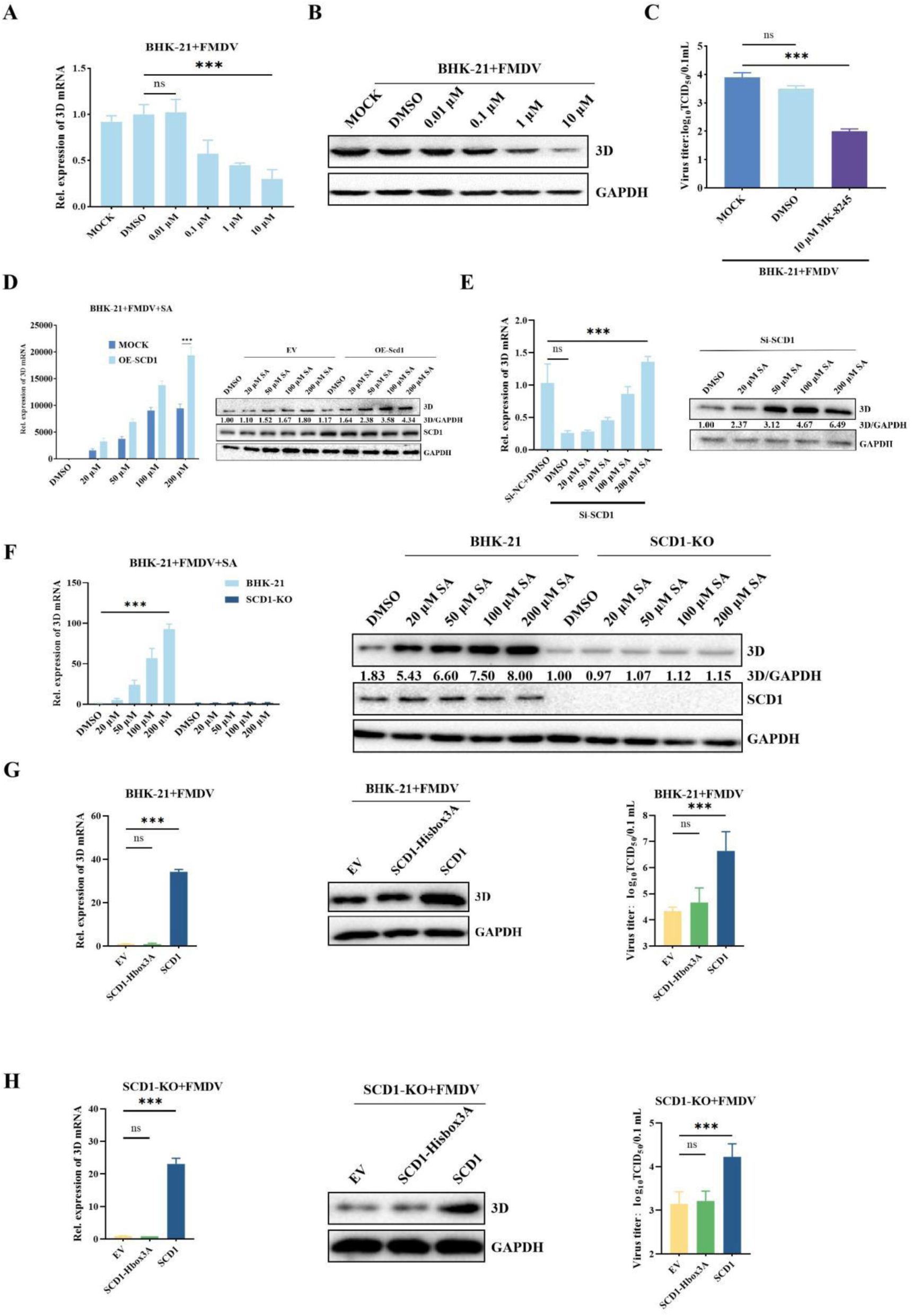
SCD1 enzyme activity is essential for promoting foot-and-mouth disease virus replication. (A) FMDV-infected BHK-21 cells were treated with maintenance medium containing different concentrations of MK8245 for 1 h, after which the samples were collected after 16 h of culture. 3D mRNA was analyzed by qPCR. (B) The 3D expression levels in A were analyzed by western blotting. (C) The viral titer in A was also determined by the TCID_50_ method. (D) FMDV-infected BHK-21 cells were treated with stearic acid (SA), and RNA was extracted and reverse-transcribed to cDNA at 16 h post infection. BHK-21 cells were transfected with OE-*SCD1*, infected with FMDV and treated with SA at 24 hpt. After 16 h of culture, the cells were harvested for 3D mRNA and protein detection. (E) BHK-21 cells were transfected with Si-*SCD1*, infected with FMDV and treated with different concentrations of SA at 24 hpt. Cells were collected at 16 hpi for 3D mRNA and protein detection. (F) *SCD1*-KO BHK-21 cells or BHK-21 cells infected with FMDV were cultured in media containing different concentrations of stearic acid, after which the cells were collected for 3D expression analysis with GAPDH normalization. (G) The effect of overexpressing wild-type SCD1 or its mutant on 3D expression. BHK-21 cells were transfected with the indicated plasmids and infected with FMDV at 16 hours post transfection. Cells were collected for 3D mRNA and protein expression analysis. The culture supernatant was harvested for virus titration using the TCID_50_ method. (H) The effect of overexpressing wild-type SCD1 or its mutant on 3D expression. *SCD1*-KO cells were transfected with the indicated plasmids and infected with FMDV at 16 hpt. Cells were collected for 3D mRNA and protein expression analysis. The culture supernatant was harvested for virus titration using the TCID_50_ method. The protein bands were quantified in grayscale by using Image-Pro Plus 6.0 software for the western blot. *n* = 3 for each group of experiments, and three parallel samples were combined for western blotting. *, *P* < 0.05; n.s., not significant.

The histidine cassette of SCD1 is involved in catalyzing the conversion of SA into unsaturated fatty acids ^[22, 23]^. To further confirm the role of SCD1 catalytic activity in regulating FMDV replication, we mutated the histidine residues at positions 297, 300, and 301 in SCD1 to alanine, to create the SCD1-Hbox3A mutant. We transfected these mutants into BHK-21 and SCD1-KO cells. Compared with those in cells transfected with SCD1, the FMDV 3D protein level was significantly lower in cells transfected with SCD1-Hbox3A (Figure 2G&H), indicating that the SCD1 histidine residues are essential for its enzymatic activity.

### SCD1 catalytic product significantly upregulates FMDV replication in BHK-Op cells and triggers FMDV replication in BHK-VEC cells

To investigate whether the SCD1-mediated inhibition of FMDV infection is dependent on unsaturated fatty acids, we examined the relationship between FMDV infection and exogenous OA. Using a CCK-8 assay, we first verified that OA is not toxic to BHK-21 cells (data not shown). Cells infected with FMDV and supplemented with different concentrations of OA were collected after 16 h of culture. Data from RT‒qPCR and western blotting analyses showed that FMDV 3D protein expression was significantly lower in *SCD1*-KO BHK-21 cells than in wild-type BHK-21 cells. OA reversed the inhibition of FMDV replication caused by knockdown of *SCD1* in a dose-dependent manner (Figure 3A). When the concentration of OA reached 200 μM, the 3D protein level was similar to that in the control group. Thus, 200 μM OA was used in subsequent experiments. The addition of 200 μM OA to FMDV-infected BHK-21 cells inhibited SCD1 enzyme activity, and the 3D protein level was similar to that in the control group (Figure 3B). FMDV replication in *SCD1*-KO cells was also increased by the addition of OA (Figure 3C). However, even when OA was added at a concentration of 200 μM, FMDV replication did not revert to normal levels.

**Figure 3.**
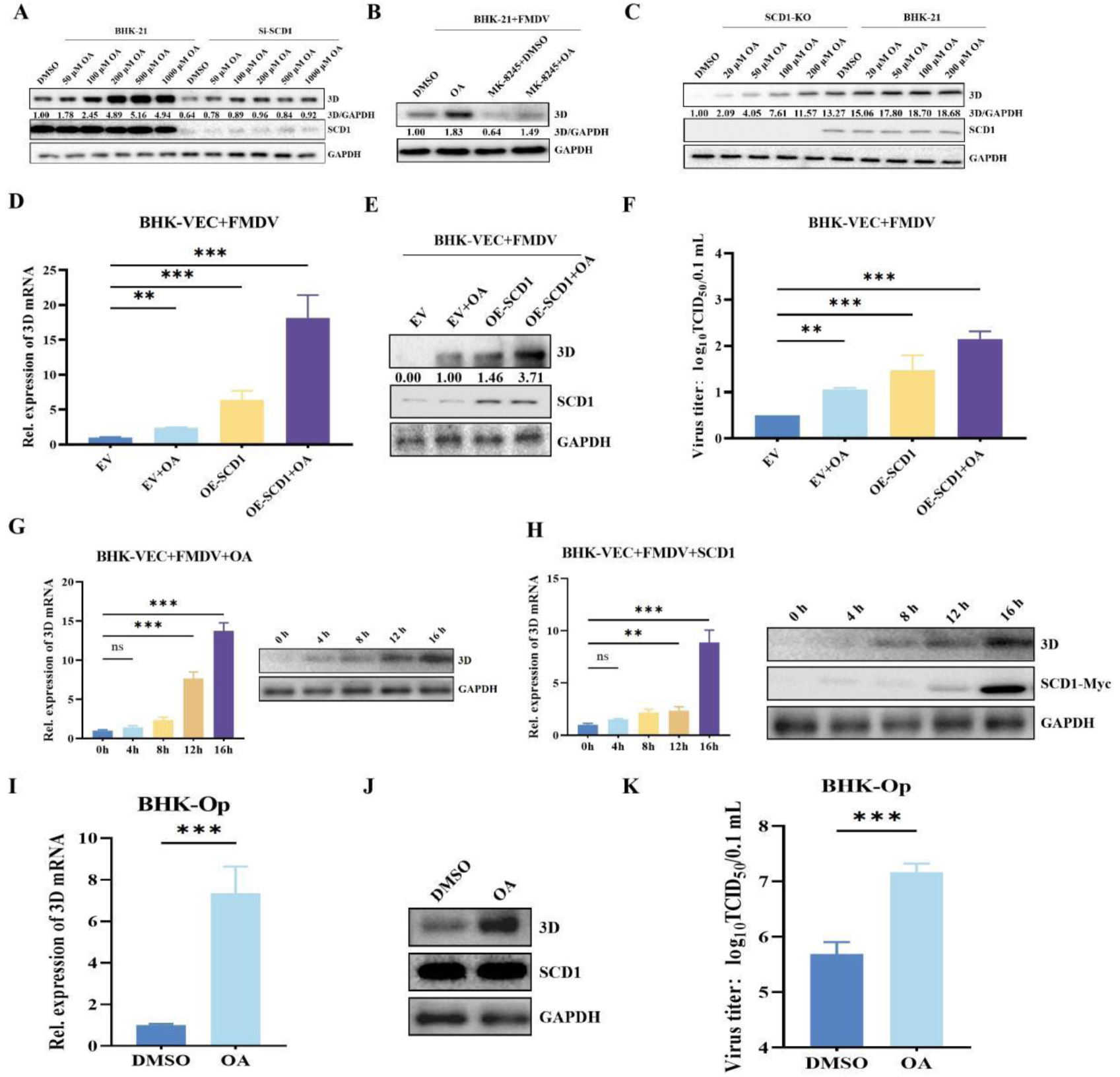
SCD1 enhances foot-and-mouth disease virus (FMDV) replicative efficiency in persistently-infected cells and triggers FMDV replication in BHK-VEC cells. (A) BHK-21 cells transfected with si-*SCD1* were infected with FMDV. At 24 hours post infection (hpi), the cells were cultured in media supplemented with different concentrations of oleic acid (OA). After 16 h of culture, the cells were harvested for 3D expression analysis. Image-Pro Plus 6.0 software was used to quantify the protein bands in the western blot. (B) FMDV-infected BHK-21 cells were treated with 20 μM MK8245 and 200 μM OA. After 16 h of culture, the cells were collected for 3D protein expression analysis. (C) *SCD1*-KO BHK-21 cells or BHK-21 cells were infected with FMDV and cultured in media containing different concentrations of OA. After 16 h of culture, the cells were harvested for 3D detection as described for GAPDH normalization. (D) BHK-VEC cells transfected with OE-*SCD1* were infected with FMDV at 16 hpt. Then, the infected cells were cultured in media supplemented with OA. After 16h of culture, the cells were collected for 3D mRNA detection. (E) The samples in Figure D were collected for 3D protein detection. (F) The viral titers were measured by the TCID50 method after cell treatment, as shown in Figure D. (G) BHK-VEC cells were infected with FMDV and cultured in media containing OA for 16 h. Cells were collected at 0, 4, 8, 12, and 16 h post infection for 3D mRNA and protein expression analysis. (H) BHK-VEC cells were transfected with Myc-*SCD1*. At 16 hpt, the cells were infected with FMDV and cultured in media containing OA for 16 h. The cells were collected at 0, 4, 8, 12, and 16 h post infection for 3D and SCD1 mRNA and protein expression analysis. (I) The BHK-Op cells were cultured in media supplemented with OA for 16 h, after which the cells were collected for 3D mRNA expression analysis. (J) The BHK-Op cells were cultured in media supplemented with OA for 16 h, after which the cells were collected for 3D protein expression analysis. (K) The viral titers were measured by the TCID50 method after cell treatment, as shown in Figure D. The protein bands were quantified in grayscale by using Image-Pro Plus 6.0 software for western blotting. *n* = 3 for each group of experiments, and three parallel samples were combined for western blotting. \**P* < 0.05, n.s., not significant.

We transfected BHK-VEC cells with the SCD1 overexpression plasmid, and at 24 hpt we infected the transfected cells with FMDV. The cells were harvested at 16 hpi for expression analysis. The data showed that overexpression of *SCD1* restored FMDV replication in BHK-VEC cells (Figure 3D-F). Exogenous addition of OA also restored FMDV replication in BHK-VEC cells. To investigate the role of *SCD1* overexpression or exogenous OA in BHK-VEC cells, treated cell cultures were collected 0, 4, 8, 12 and 16 h after FMDV infection and analyzed by RT‒qPCR and western blotting. In the OA-treated group, FMDV 3D protein expression was most significantly upregulated at 12 hpi, while in the SCD1-overexpressing group, FMDV 3D protein expression was most significantly upregulated at 16 hpi (Figure 3G and H).

We further investigated the regulatory effect of exogenous OA on FMDV replication in BHK-Op cells. The data showed that FMDV 3D expression and FMDV titer were significantly increased in BHK-Op cells after exogenous OA addition (Figure 3I-K). These results indicate that SCD1 regulates FMDV replication in BHK-Op cells and activates FMDV replication in BHK-VEC cells.

### SCD1 recruits FMDV 2C to detergent-resistant membranes to promote FMDV replication complex establishment

To understand the functional mechanism that SCD1 mediates FMDV replication complex formation, we further investigated whether SCD1 interacts with the nonstructural 2C protein located on the FMDV replication complex. The Flag-tagged FMDV 2C and Myc-tagged SCD1 were cotransfected into BHK-21 cells. The cells were infected with FMDV at 24 hpt, and the infected cells were collected at 16 hpt. IP data (Figure 4A) showed that SCD1 interacted with 2C, an FMDV replication complex protein that plays a key role in both membrane rearrangement and formation of the viral replication complex^[24]^. Further study revealed that in FMDV replication complexes containing FMDV 2C, vimentin and SCD1. Co-IP experiments (Figure 4B) confirmed the interactions among the three proteins.

**Figure 4.**
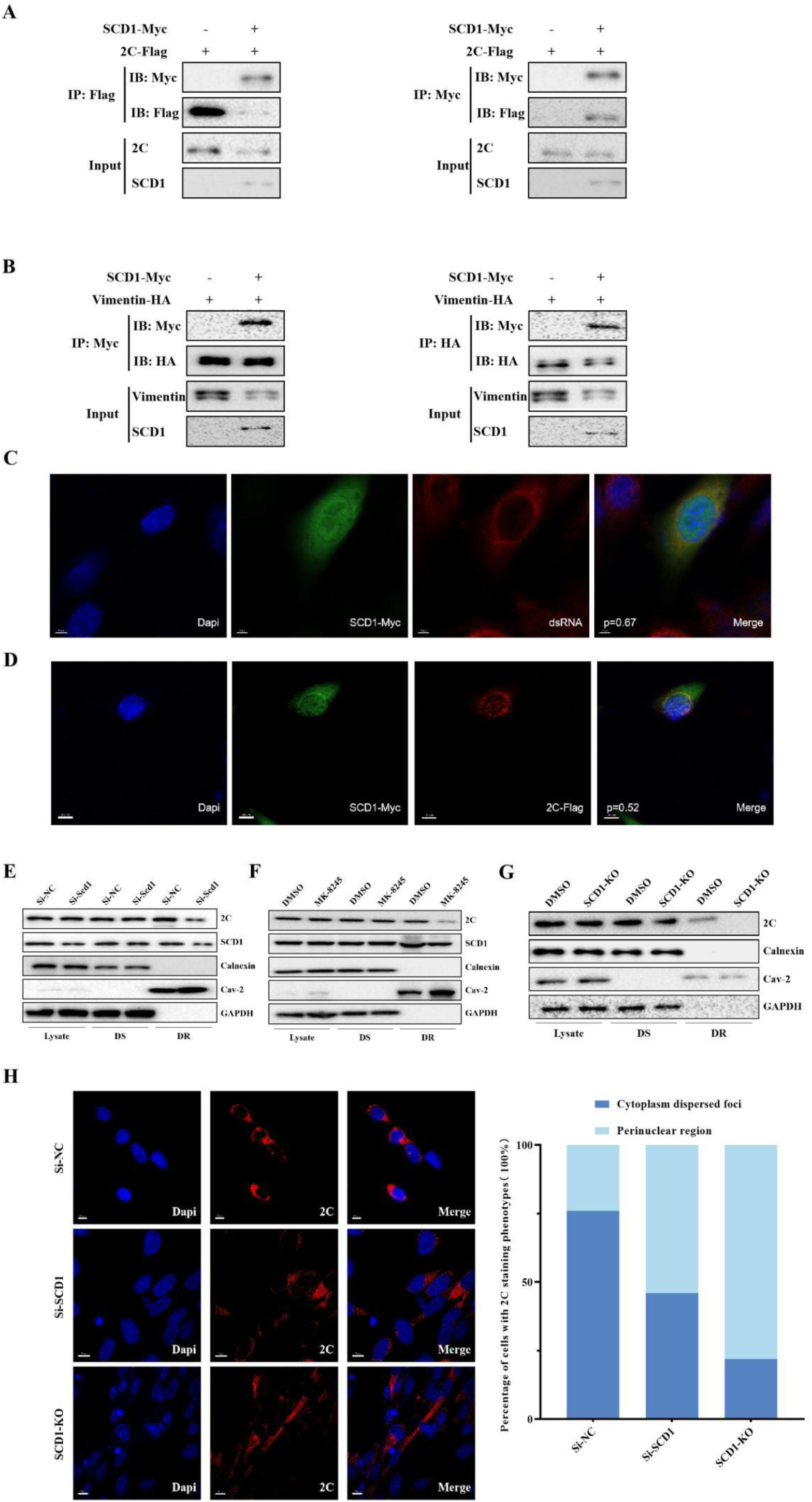
SCD1 recruits foot-and-mouth disease virus (FMDV) 2C to detergent-resistant membranes. (A) BHK-21 cells were cotransfected with SCD1-Myc and FMDV 2C and a Flag-tag expression plasmid (Flag-2C). The cells were harvested at 24 hours post transfection (hpt) to detect the interaction between SCD1 and 2C by coimmunoprecipitation (co-IP) assays; (B) BHK-21 cells were cotransfected with SCD1-Myc and HA-tagged vimentin (Vimentin-HA). The cells were collected at 24 hpt to detect the interaction between SCD1 and vimentin by co-IP assays. (C) BHK-21 cells were transfected with SCD1-Myc, and the cells were infected with FMDV at 24 hpt. After 6 h of culture, the localization of SCD1 and dsRNA was analyzed using anti-SCD1 or dsRNA antibody staining. DAPI staining was used to label the nuclei (blue). (D) BHK-21 cells were cotransfected with SCD1-Myc and Flag-2C, and the cells were infected with FMDV 24 hpt. After 6 h of culture, the localization of SCD1 and 2C was analyzed using antibodies directed against SCD1 and 2C by IFA. DAPI staining was used to label the nucleus (blue). The Pearson coefficients for C and D are labeled on the corresponding merged plots. (E) BHK-21 cells were transfected with *SCD1*-KO cells, and the cells were infected with FMDV at 24 hpt. After 16 h of culture, the cells were collected and lysed, and the lysates were separated into detergent-soluble or detergent-insoluble membrane fractions. Cell lysates and detergent-soluble (DS) and detergent-resistant (DR) membrane fractions were analyzed by immunoblotting with the indicated antibodies. (F) BHK-21 cells were infected with FMDV and cultured in medium supplemented with an SCD1 inhibitor. After 16 h of culture, the cell lysates, DSs, and DR membrane fractions were analyzed. (G) *SCD1*-KO cells were infected with FMDV. At 16 hpi, the cells were harvested and lysed, and the cell lysates, DS, and DR membrane fractions were analyzed. (H) Localization of FMDV 2C in *SCD1*-knockdown BHK-21 cells and *SCD1*-KO BHK-21 cells. For *SCD1* knockdown, cells were transfected with si-*SCD1*. At 24 hpt, the cells were infected with FMDV. After 16 h of culture, the subcellular localization of 2C was analyzed using an anti-2C antibody and IFA. DAPI staining was used to label the nuclei (blue; upper panel). Fifty 2C-positive cells were randomly selected for counting (expressed as a percentage). Three independent replicates were performed for each set of experiments.

To further explore whether SCD1 is located within the FMDV replication complex, we used IF to probe whether SCD1 colocalizes with the replication complex marker dsRNA. We transfected an Myc-SCD1 plasmid into BHK-21 cells, and the transfected cells were infected with FMDV at 24 hpt. We determined colocalization of SCD1, 2C, and dsRNA components based on the Pearson coefficient. If the Pearson coefficient was >0.5, the two components were considered to be colocalized. The data showed that SCD1 colocalized with dsRNA, and that 2C had the same localization pattern as SCD1 (Figure 4C&D), suggesting that SCD1 is a constituent protein of the FMDV replication complex.

The classical approach to studying plasma membrane microregions involves obtaining detergent-resistant extracted membrane fractions from cell membranes and analyzing their composition. In HCV, the proteins involved in replication complexes have been shown to be highly detergent resistant^[6]^, suggesting that the synthesis of HCV RNA occurs on lipid raft membrane structures. HCV nonstructural proteins exhibit strong detergent resistance and are present on the lipid rafts of nonstructural protein NS3-5B-overexpressing cells^[25]^. Therefore, probing the membrane localization and detergent resistance of the FMDV nonstructural 2C protein is highly important for investigating the detailed mechanism through which 2C-mediates host cell membrane rearrangement.

We lysed *SCD1*-KO BHK-21 cells and SCD1 inhibitor-treated BHK-21 cells. Then, the cells were collected and divided into detergent-resistant and detergent-sensitive membrane fractions. Knockdown of *SCD1* significantly downregulated 2C in detergent-resistant membranes. Similarly, significantly down-regulated 2C concentration was observed in the detergent-resistant membranes of inhibitor-treated cells (Figure 4E, F and G). In the inhibitor-treated group, the 2C concentration in the detergent-resistant membrane was also significantly reduced, probably due to the inhibition of SCD1 activity and consequent inability to recruit 2C.

To further investigate the membrane rearrangement induced by the 2C-mediated recruitment of SCD1, we observed the subcellular localization of 2C in *SCD1*-KO cells. After FMDV infection, 2C formed large and concentrated foci around the nucleus, suggesting that 2C was enriched in the region where replication complexes are concentrated (Figure 4H). The distribution and subcellular localization of 2C in *SCD1*-KO cells revealed a dispersed pattern and increased disorder (Figure 4H), suggests that the recruitment of SCD1 to 2C is dose-dependent. The data showed that SCD1 loses its ability to recruit 2C when it loses the ability to interact with 2C. Statistical analysis of data from 100 randomly selected cells (Figure 4H) revealed that SCD1 plays an important role in the recruitment of 2C, while the 2C distribution in cells became scattered after decreasing the SCD1 concentration, with an observed decrease in the 2C protein in the perinuclear region. In conclusion, knockdown or knockout of *SCD1* altered the subcellular localization of 2C. Complementation with SCD1 mutants that cannot interact with 2C failed to change the localization of 2C. In contrast, in the FDMV-infected SCD1 KO cell line, 2C localization was disorganized, while FMDV replication was attenuated. We speculated that proper localization of 2C to the FMDV replication complex facilitates FMDV replication. The data indicate that in infected cells 2C is mainly localized to the perinuclear region and is partially scattered in the cytoplasm, and that this distribution is determined by the ability of SCD1 to interact with 2C.

### SCD1 and its enzymatic products are involved in FMDV replication complex formation

To further investigate the role of SCD1 in the FMDV replication complex, we observed replication complexes in BHK-21, BHK-VEC, and BHK-Op cells by TEM. We also observed the number of replication complexes and their morphology at overexpressed or knocked down *SCD1* in BHK-21 and BHK-VEC cells to study the effect of SCD1 on the establishment of replication complexes. Both the number of replication complexes and lipid droplets were greatly reduced in cells with SCD1 knockdown compared to cells without SCD1 knockdown (Figure 5A, G). After the addition of exogenous OA, the number of lipid droplets and FMDV replication complexes in the cells significantly increased (Figure 5B, G). This difference was more pronounced in BHK-Op cells than in the other cell lines. In BHK-Op cells, the endoplasmic reticulum was deformed and usually connected to mitochondria, suggesting that the endoplasmic reticulum requires additional energy to maintain its normal morphology or to navigate to form replication complexes in these cells. When SCD1 was overexpressed, the number of replication complexes and lipid droplets significantly increased (Figure 5C, G).

**Figure 5.**
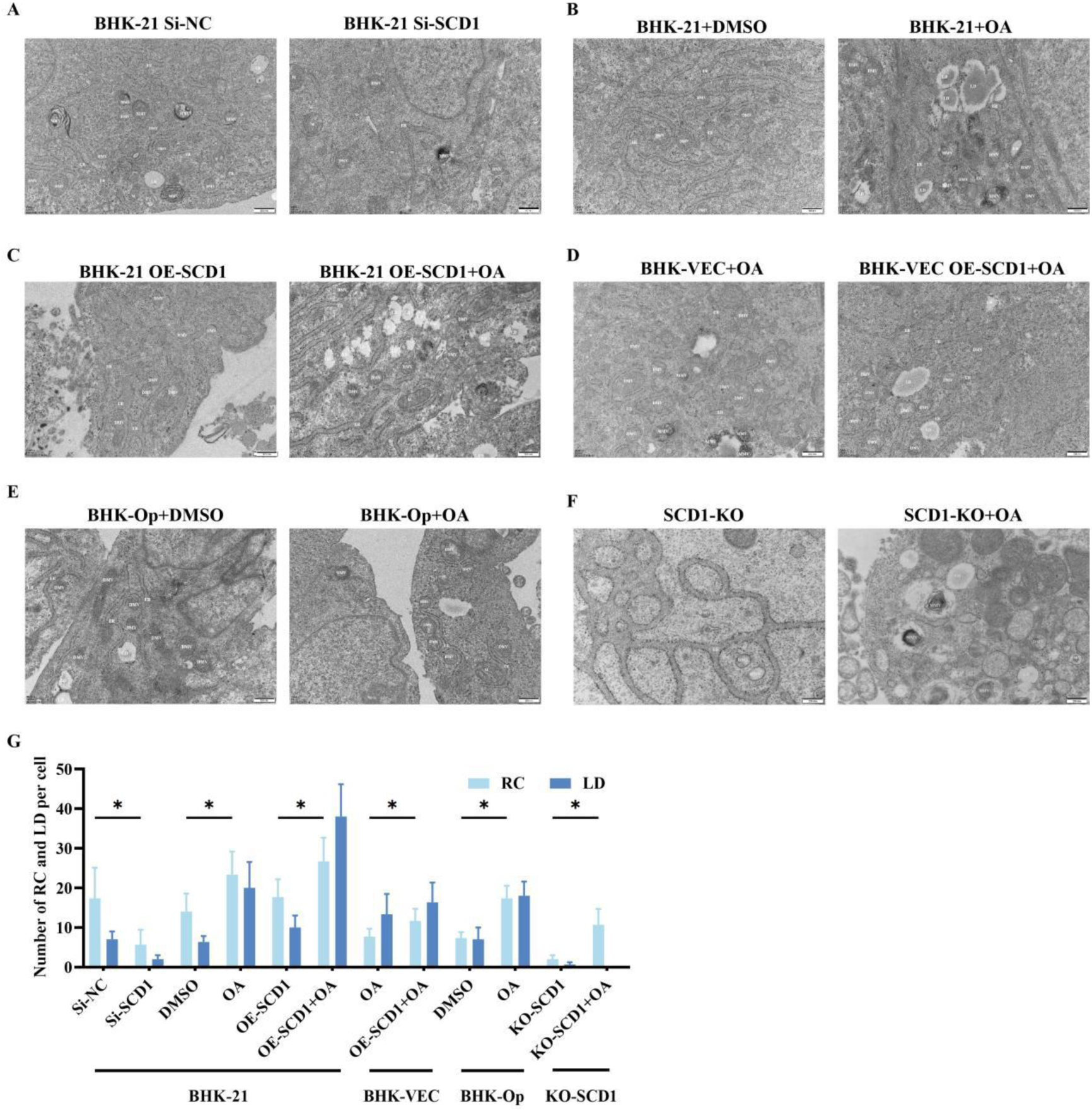
SCD1 and its enzyme products are involved in foot-and-mouth disease virus (FMDV) replication complex formation. (A–F) Transmission electron microscopy was performed as described in the “Methods” section. In the figures, “M” indicates mitochondria, “ER” indicates the endoplasmic reticulum, DMV indicates “double-membrane vesicles”, and LD indicates “lipid droplets”. Scale bars, 200 nm and 500 nm. (G) The number of RCs and LDs in five random cells.

We did not observe replication complexes in FMDV-infected BHK-VEC cells (data not shown), which may explain why FMDV cannot replicate in BHK-VEC cells. However, after the addition of exogenous OA or expression of SCD1, replication complexes and lipid droplets formed in BHK-VEC cells (Figure 5D, G). Multiple lamellar vesicles (MMVs) were produced at the same time, but the number of MMVs was markedly lower and the MMVs were more dispersed than those in BHK-21 cells, which was consistent with our previous finding that overexpression of SCD1 or exogenous OA could not completely restore FMDV replication in BHK-VEC cells. The addition of OA to BHK-Op cells resulted in increased replication complexes and lipid droplets (Figure 5E). Finally, the addition of exogenous OA to SCD1-KO cells increased the number of intracellular lipid droplets and replication complexes (Figure 5F, G). These results further demonstrated that upregulation of SCD1 activity led to endoplasmic reticulum rupture in host cells, which in turn promoted FMDV replication complex formation. In conclusion, upregulation of SCD1 activity leads to an increase in the number of replication complexes and lipid droplets. These findings suggest that OA is a necessary raw material for the formation of lipid droplets and replication complexes in cells. In contrast, down-regulation of SCD1 activity decreases the number of replication complexes and lipid droplets and also leads to abnormal FMDV replication complex morphology.

### The replication of a broad spectrum of positive-sense RNA viruses is regulated by SCD1 enzyme activity

To further explore whether SCD1 regulates FMDV replication in cells from other species, we knocked down *SCD1* expression using RNAi or inhibited its enzyme activity using SCD1 inhibitors in PK-15 cells (a pig kidney epithelial cell line) and observed a significant decrease in FMDV replication efficiency (Figure 6A&B). To evaluate the effect of SCD1-regulated lipid metabolism in on the replication of other positive-sense RNA viruses, we used the positive-strand RNA viruses PV1, REO176, and EV71 in further experiments. First, we detected the RNA levels of PV1, REO176 and EV71 in Vero cells transfected with si-SCD1 by RT‒qPCR. REO176 replication was inhibited in si-SCD1-transfected cells (Figure 6C). After adding an appropriate amount of inhibitor to inhibit the activity of SCD1, we added 200 μM of OA. the results showed that the decrease in the replication level of REO176 was reversed(Figure 6C). This result was further confirmed by measuring TCID_50_ of viral titers (Figure 6D). We then added different concentrations of the SCD1 inhibitor to Vero cells, followed by supplementation with OA. Consistent with the *SCD1* knockdown, the exogenous SCD1 inhibitor inhibited REO176 replication, while supplementation with OA reversed this inhibition (Figure 6E). Inhibition of SCD1 activity decreased REO176 titers, whereas supplementation with OA upregulated REO176 titers (Figure 6F). Similar results were observed for other positive-sense RNA viruses, PV1 (Figure 6G-J), and EV71 (Figure 6K-N). Thus, SCD1 is important for the replication of multiple positive-sense RNA viruses in a variety of mammalian cells.

**Figure 6.**
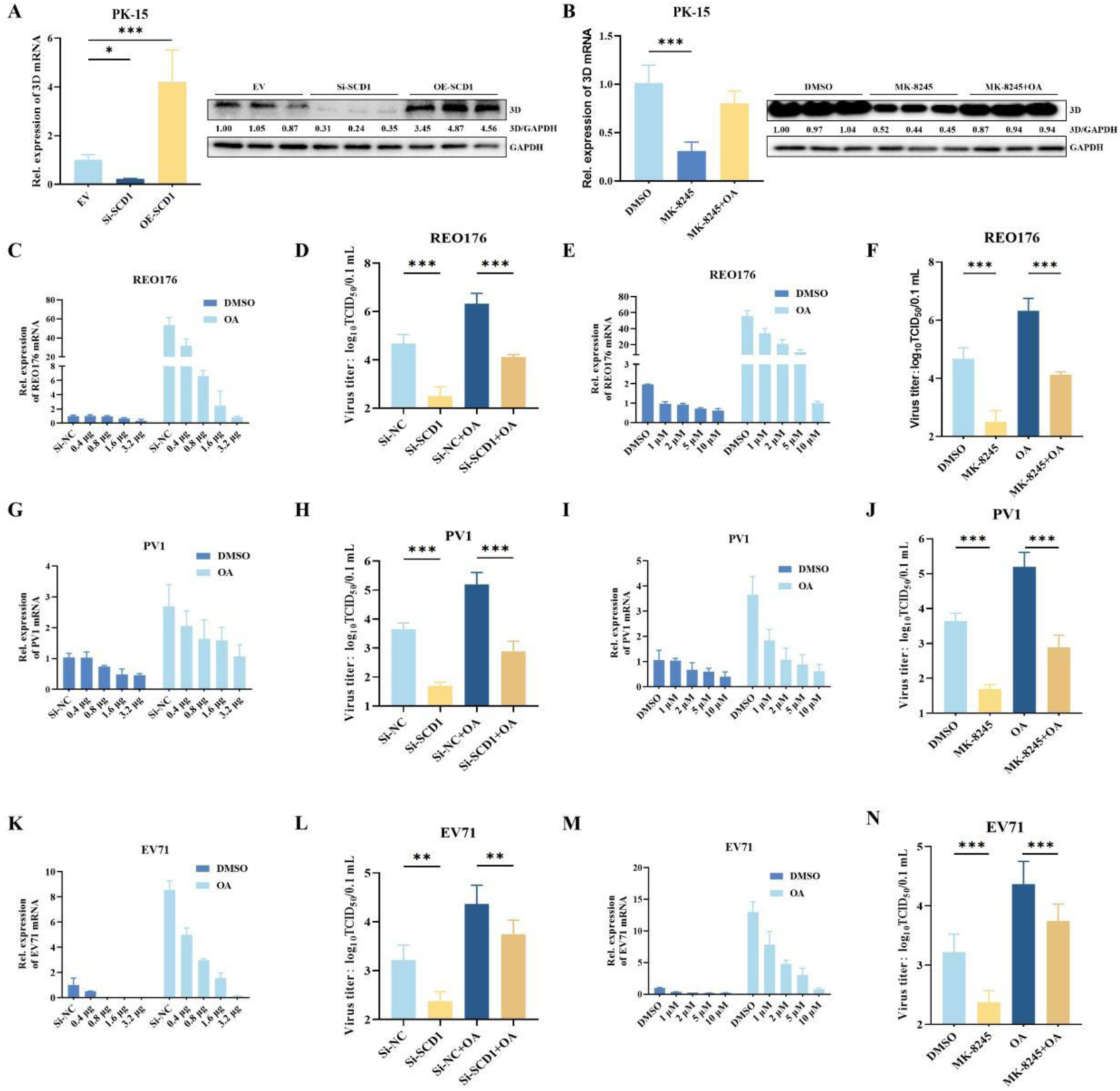
A broad spectrum of positive-strand RNA virus replication is regulated by SCD1 enzyme activity. (A) PK-15 cells were transfected with OE-SCD1 or Si-*SCD*, and at 24 hours post transfection (hpt), the cells were infected with FMDV. After 16 h of culture, the expression of FMDV 3D was detected by qPCR and western blotting. (B) FMDV-infected PK-15 cells were cultured in media supplemented with an SCD1 inhibitor with or without OA. Cells were collected at 16 hours post infection (hpi) for 3D expression analysis. (C & D) The medium of REO176-infected Vero cells was cultured in 2% minimal essential medium (MEM) for 3 h. The cells were collected after an additional 16 h of culture for vRNA detection (C); the culture supernatant was collected for virus titration by the TCID_50_ method (D); (E & F) REO176-infected Vero cells were cultured in medium containing different concentrations of MK8245, and the cells were collected at 16 hpi for vRNA detection (E); or the culture supernatant was collected for virus titration (F); (G) PV1-infected Vero cells were cultured in media containing different concentrations of OA. After 16 h of culture, the cells were collected for vRNA detection. (H) PV1-infected Vero cells were cultured in media containing MK-8245, OA or MK-8245+ OA. After 16 h of culture, the culture supernatant was collected for virus titration. (I) Vero cells were infected with PV1, and after 3 h, the medium was replaced with maintenance medium containing different concentrations of MK8245. After 16 h, RNA was extracted from the cells and reverse transcribed into cDNA, and the RNA level of PV1 was detected by qPCR. (J) Vero cells were infected with PV1, and after 3 h, the medium was replaced with maintenance medium containing different concentrations of MK8245. The supernatant was collected after an additional 16 h of culture, and the viral titer in the supernatant was determined by the TCID_50_ method. (K) EV71-infected Vero cells were cultured in media containing different concentrations of OA. After 16 h of culture, the cells were collected for virus RNA detection. (L) EV71-infected Vero cells were cultured in media containing MK-8245, OA or MK-8245+ OA. After 16 h of culture, the supernatant was collected for virus titration; (M) Vero cells were infected with EV71 for 3 h, after which the medium was replaced with maintenance medium containing different concentrations of MK8245. After 16 h of culture, the cells were collected for EV71 RNA detection by qPCR. (N) Vero cells were infected with EV71, and after 3 h, the medium was replaced with maintenance medium containing different concentrations of MK8245. The supernatant was collected 16 h later, and the virus titer was determined via the TCID_50_ method.

## Discussion

FMDV replication is closely related to host cell lipid metabolism, and SCD1 is the rate-limiting enzyme that catalyzes production of the monounsaturated fatty acid OA from stearic acid. By regulating SCD1 expression, we were able to influence FMDV replication in BHK-21, BHK-VEC and BHK-Op cells. The addition of either the catalytic product of SCD1 or its substrate, stearic acid, to these cells promoted FMDV replication. Interestingly, OA addition or SCD1 overexpression in BHK-VEC cells, in which SCD1 expression is usually low, resulted in FMDV replication, . When we exogenously added moderate amounts of OA to FMDV-infected cells in which SCD1 could be knocked down or knocked out, FMDV replication was restored to normal levels. Our results also showed that the SCD1 enzymatic activity, to produce OA, is controlled by the AMP-activated protein kinase (AMPK) pathway and that promotion of AMPK pathway signaling inhibited FMDV replication (Figure S2). Although the SCD1 gene, an important host factor affecting FMDV replication, was involved in the establishment of the BHK-Op and BHK-VEC cell models, the establishment of the model was the result of the coordination of multiple genes.

To investigate the effect of SCD1 catalytic activity on FMDV replication, we constructed a histidine point mutant of SCD1 and found that inactivation of SCD1 enzyme activity led to significant downregulation of FMDV replication. The binding of SCD1 to 2C may play a role in assisting FMDV replication, but the exact mechanism through which this occurs is still unclear. IF experiments confirmed the colocalization of SCD1 with 2C in the replication complex. Interestingly, SCD1, an endoplasmic reticulum-anchored protein, is not normally found in other subcellular structures. However, after infection with FMDV, the host cell undergoes membrane rearrangement, and this membrane rearrangement likely leads to SCD1 entry into the bilayer of the replication complex and its colocalizatoin with 2C. Furthermore, 2C is both involved in and mediates membrane rearrangement in FMDV host cells, which correlates with our previous finding that OA is involved in the synthesis of membrane structures with negative curvature ^[26, 27]^. In addition, functional studies of 2C proteins suggest that early membrane rearrangements may be an important process for subsequent bilayer vesicle formation. Our results suggest that SCD1 recruits 2C and plays a role in providing lipids for vesicle synthesis and that the OA produced by SCD1 is involved in cell membrane formation. This process is necessary for the establishment of the FMDV replication complex. However, the detailed mechanism through which SCD1 functions in this context needs to be further investigated.

We extracted detergent-resistant membrane structures from cells and found that the level of 2C in detergent-resistant membranes was significantly lower in *SCD1*-knockdown cells than in normal BHK-21 cells. In addition, we performed IF experiments and found that *SCD1* knockdown shifted the subcellular localization of 2C from the perinuclear region, where the FMDV replication complex is located, to the cytoplasm. These results suggest that the decrease in SCD1 activity leads to a change in 2C localization and a decrease in the amount of 2C in the detergent-resistant membranes of the replication complexes. We suggest that knockdown of *SCD1* inhibits FMDV replication, likely not only by decreasing the enzymatic activity of SCD1 and decreasing OA production, which in turn leads to decreased replication complexes but also through its involvement in recruiting nonstructural FMDV proteins, which localize 2C to the replication complex and function in 2C-mediated membrane rearrangement, facilitating the establishment of the FMDV replication complex. Additionally, our results suggest that the replication complex contains a detergent-resistant membrane structure, and the proteins associated with FMDV replication may all be located within this structure. However, it is worth noting that both our previous demonstration of the role of SCD1 enzymatic activity and validation of the 2C recruitment function of SCD1 revealed that there are other factors in host cells that can recruit nonstructural FMDV proteins and that SCD1 is not the only host factor that is important for the establishment of the FMDV replication complex. The discovery of other host factors with similar abilities to recruit nonstructural proteins or promote the establishment of the FMDV replication complex is one of the main directions of our future work.

A common feature of positive-strand RNA viruses is that they cause remodeling of the inner membrane of infected cells. To undergo replication, positive-strand RNA requires a membrane spacer, in which dsRNA, a marker of the replication complex, is produced before viral assembly^[28]^. The membrane source of bilayer vesicles, the replication complex, may also contain other membrane structures, such as lipid droplets, mitochondria, and early and late endosomes^[29]^. Our results indicated that the replication complex is located near the endoplasmic reticulum, and we observed multilayered vesicles and bilayered vesicles. Such multilayered vesicles are thought to occur in the late stages of HCV infection and are not associated with RNA replication. In addition, the inhibition of SCD1 activity resulted in structural abnormalities and a decrease in the number of replication complexes. Conversely, overexpression of SCD1 or the addition of OA increased the number of replication complexes, thus facilitating viral replication. Notably, we successfully reconstructed the replication complex in BHK-VEC cells by upregulating SCD1 activity, suggesting that abnormal formation of the replication complex is one of the important reasons why FMDV fails to replicate properly in BHK-VEC cells. However, the structure of the replication complex in these cells was still abnormal after SCD1 overexpression, suggesting that there are other host factors are also involved in the formation of the replication complex. The above evidence strongly suggests that SCD1 regulates host cell lipid metabolism and participates in the replication complex formation, which in turn regulates FMDV replication. Additionally, in knockout cells, SCD1 promoted the collapse of the endoplasmic reticulum after FMDV infection of host cells, thereby promoting the formation of replication complexes. This finding provides key evidence that SCD1 is directly involved in the formation of the FMDV replication complex. Taken together with the results of previous studies, these findings suggest that SCD1 is involved in recruiting FMDV 2C to the replication complex.Therefore, disruption of SCD1 activity can somewhat reduce the 2C content in the replication complex, which provides a potential target for the development of anti-FMDV drugs.

To investigate the role of SCD1-mediated lipid metabolism in promoting the replication of positive-strand RNA viruses in general, we inhibited SCD1 activity and observed the regulation of REO176, PV1, and EV71 viral replication. Inhibition of SCD1 activity downregulated the replication of REO176, PV1, and EV71, while the addition of exogenous OA reversed this inhibition. This finding suggested that SCD1 also regulates the replication of these positive-strand RNA viruses by modulating lipid metabolism and thus the formation of the viral replication complex. SCD1 is a potential target for the treatment of RNA virus infections, and our findings provide new ideas for the development of antiviral drugs.

## Figure legends

**Figure S1.**
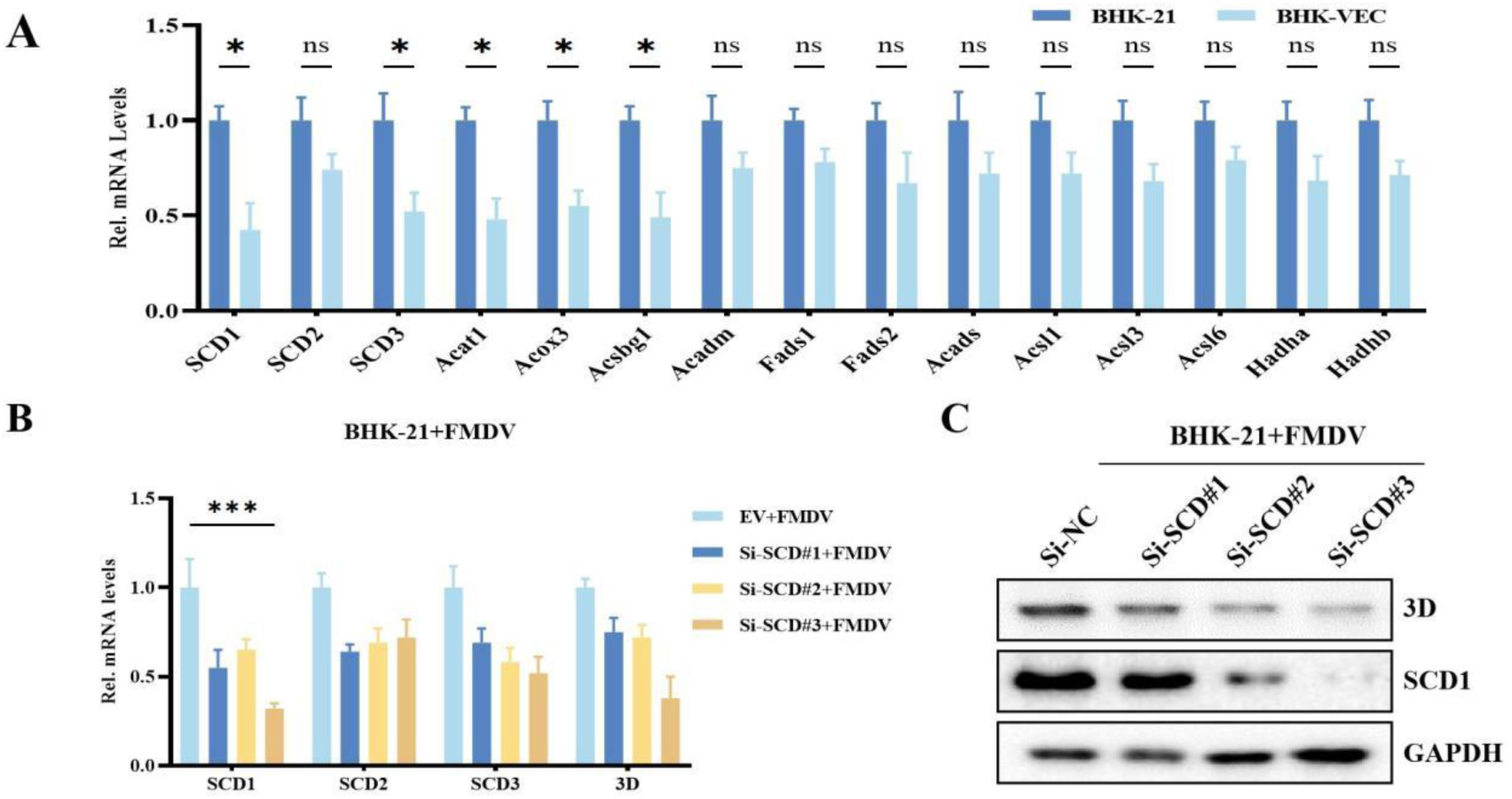
Functional characterization of the *SCD1* gene. (A) Expression analysis of genes enriched in the fatty acid metabolism pathway via transcriptome sequencing of BHK-VEC cells and BHK-21 cells; (B) BHK-21 cells transfected with the SCD gene RNAi vector were infected with foot-and-mouth disease virus (FMDV) at 16 hours post transfection (hpt). After 24 h of culture, the infected cells were harvested for SCD gene or 3D detection via qPCR. (C) BHK-21 cells transfected with the SCD gene RNAi vector were transfected with FMDV at 16 (hpt). After 24 h of culture, the infected cells were collected and subjected to immunoblotting to detect the protein levels of 3D and SCD1.

**Figure S2.**
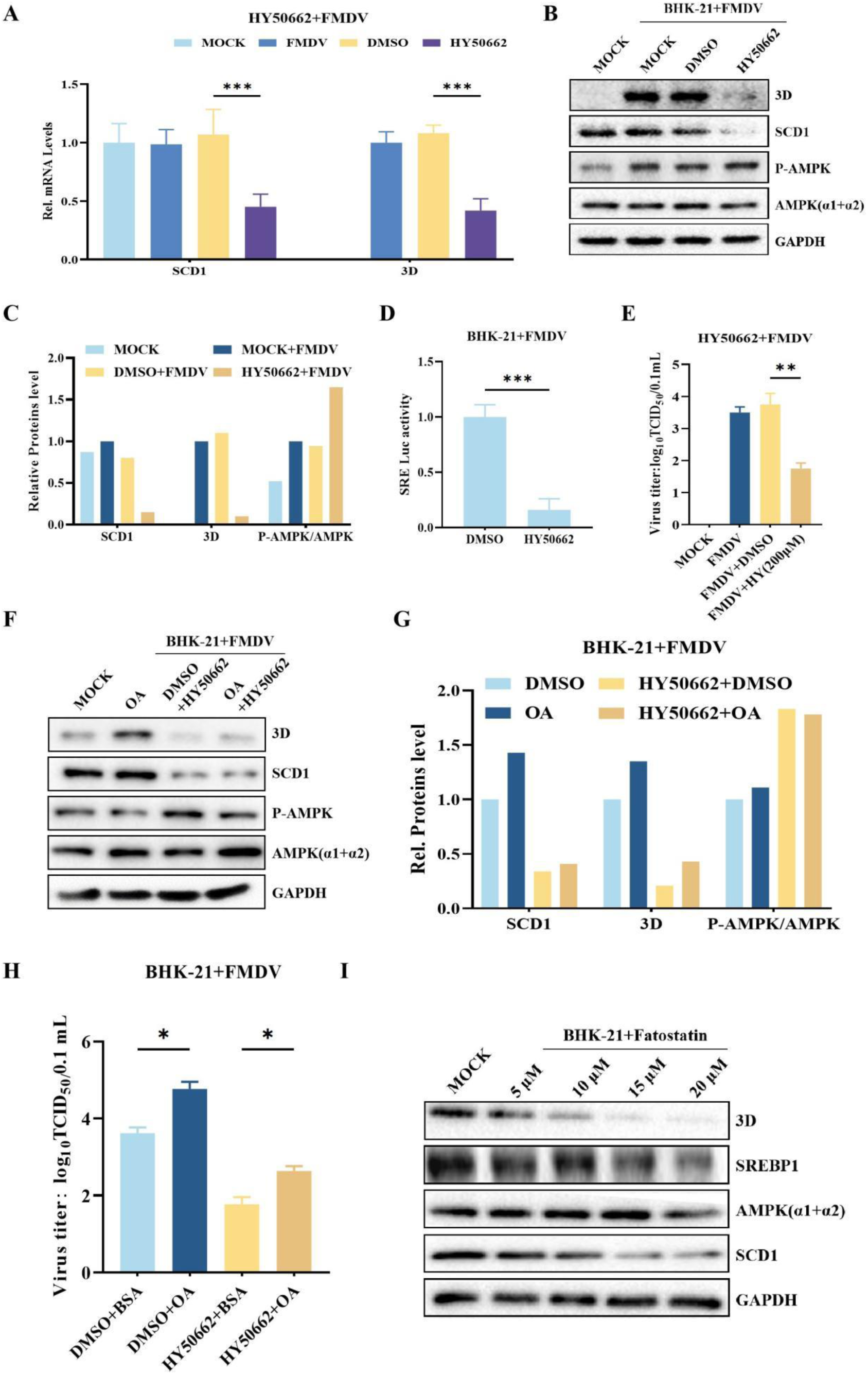
SCD1-mediated regulation of FMDV replication is dependent on the AMPK pathway. (A) FMDV-infected BHK-21 cells were cultured in media supplemented with HY50662 for 16 h, and infected cells were collected for detection of SCD1 or 3D mRNA levels via qPCR or (B) for detection of AMPK, p-AMPK, SCD1, or 3D protein. (C) Image-pro Plus 6.0 software was used to quantify the gray values of the protein bands from the western blot in Figure C, after which the gray values of the target proteins were compared to those of the corresponding internal references. (D) FMDV-infected BHK-21 cells were cultured in medium containing HY50662. After 16 h of culture, the cells were collected and lysed for transcriptional activation level detection using a luciferase reporter or (E) for determination of the viral titer in the supernatant using the TCID50 method. (F) FMDV-infected BHK-21 cells were cultured in medium containing HY50662, after which the BHK-21 cells were treated with different concentrations of OA. After 16 h of culture, the cells were subjected to AMPK, p-AMPK, SCD1, or 3D protein detection. (G) Image-Pro Plus 6.0 software was used to analyze the protein bands of the western blot in Figure C. The protein bands of the western blot were quantified by gray value, and the gray value of the target protein was subsequently compared to the gray value of the corresponding internal reference. (H) FMDV-infected BHK-21 cells were cultured in medium containing HY0662 and different concentrations of OA. The supernatant was collected at 16 h postinfection for viral titration using the TCID50 method. (I) FMDV-infected BHK-21 cells were cultured in Fatostain-containing medium. After 16 h of culture, the infected cells were harvested for SREBP1, AMPK, SCD1, or 3D protein detection. (n=3 for each group of experiments; *p<0.05, **p<0.01, n.s. not significantly different)

## Acknowledgments

This work was supported by the financial support of the National Science and Technology Infrastructure Grants (NSTI-BMCR21, NSTI-BMCR22, NSTI-BMCR23) to Dr. Chao Shen.

## Author Contributions

Study design, C.S. X.T. and B.L. Experiments performation, B.L. J.H. and Z.Y. Data curation. Project administration, C.S. Funding acquisition, C.S. Writing - original draft, B.L. Y.Y. Z.Y. and Y.S. Writing - review & editing, C.S. and Z.P..

